# Streaming of repeated noise in primary and secondary fields of auditory cortex

**DOI:** 10.1101/738583

**Authors:** Daniela Saderi, Brad N. Buran, Stephen V. David

## Abstract

Statistical regularities in natural sounds facilitate the perceptual segregation of auditory sources, or streams. Repetition is one cue that drives stream segregation in humans, but the neural basis of this perceptual phenomenon remains unknown. We trained ferrets to detect a stream of repeating noise samples (foreground) embedded in a stream of random noise samples (background). While they listened passively, we recorded neural activity in primary (A1) and secondary (PEG) fields of auditory cortex. We used context-dependent encoding models to test for evidence of streaming of the repeating stimulus in these brain areas. Separate models tested whether time-varying neural spike rates were better predicted by scaling the response to both streams of the repeating stimulus equally (global response gain), or by scaling the response of one stream relative to another (stream-specific response gain). Consistent with adaptation, we found an overall reduction in global gain when the stimulus was repeated. However, when we measured stream-specific changes in gain, neural responses to the foreground stream were enhanced relative to the background. This enhancement was stronger in PEG than in A1. In A1, the degree of enhancement depended on auditory tuning. It was strongest in units that displayed low sparseness (*i.e*., broad sensory tuning) and were tuned preferentially to the repeated sample. Thus, while overall auditory responses were reduced by the repeating sound, enhancement of responses to the foreground stream relative to the background provides evidence for stream segregation that emerges in A1 and is refined in PEG.

## Introduction

Sounds generated by different sources, or auditory objects, impinge on the ear as a complex mixture, with acoustic energy generated by each source overlapping in both time and frequency. The auditory system has the remarkable ability to group these dynamically changing spectro-temporal sound features into percepts of their distinct sources, in a process known as auditory streaming (1, 2). Streaming requires statistical analysis of sound sources: streams that come from the same sound source share statistical regularities, and the brain uses these properties as cues for stream integration or segregation (2–6).

Basic acoustic features, such as separation in frequency and time, are key perceptual cues for segregating simple, alternating sequences of pure tones (7–10). However, more complex natural sounds often overlap in frequency. Segregating spectrally overlapping sounds requires use of perceptual cues such as pitch (11, 12), timbre (13–15), spatial location (3, 12–16), common onset (17, 18), and temporal regularity (19–22). In a relevant study, McDermott *et al*. (2011) tested specifically for the benefit of temporal regularity with a set of naturalistic noise samples that lacked other cues for streaming (23). While non-repeating samples could not be distinguished from background noise, humans could identify these same samples when they were repeated. The neural bases of this perceptual pop-out remain unknown.

In contrast to the robust perceptual enhancement reported for a repeating foreground stream, studies of neurophysiological activity in auditory cortex have emphasized a suppressive effect of repetition (24). Single neurons undergo stimulus-specific adaptation (SSA), where responses to repeated tones adapt, but responses to an oddball stimulus, such as a tone at a different frequency, are less adapted or even facilitated, reflecting perceptual pop-out of the oddball sound (25, 26). In human electroencephalography (EEG), a possibly related phenomenon is observed in a late event-related component, called the mismatch negativity (MMN). Although the dynamics are slower than SSA, MMN is also elicited by rare deviant sounds randomly interspersed among frequent standard sounds (27). There is no evidence that links SSA or MMN with repetition-based grouping, but it is possible that these processes share some of the same neural circuits. How the brain might use adaptation to a repeating sound to enhance its perception is not known.

In this study, we investigated neural correlates of streaming induced by repetition of complex sounds in primary (A1) and secondary (PEG) fields of the auditory cortex. We first established the ferret as an animal model for streaming of repeating noise sounds, by designing a behavioral paradigm that assessed animals’ ability to detect repetitions embedded in mixtures. We then recorded single- and multi-unit activity in A1 and PEG of un-anesthetized, passively listening ferrets, either trained or naïve to the detection task. We tested the prediction that auditory cortical neurons facilitate stream segregation by selectively enhancing their response to the repeating (*i.e*., foreground) stream. We used context-dependent sound encoding models to quantify the relative contribution of the two overlapping streams to the evoked neural response (28). We found that neural responses to the repeated stimuli were reduced overall in both areas, consistent with previous studies that reported adaptation for a single repeating stream (25). Additionally, some neurons in both cortical fields displayed foreground-specific responses that were enhanced with respect to responses to the simultaneous background stream. These results provide evidence for a model of streaming cued by repetition that starts in primary and is refined in secondary fields of the auditory cortex.

## Results

### Ferrets perceive repeated patterns embedded in noise

To investigate the physiological underpinnings of repetition-based streaming in an animal model, we first developed a behavioral paradigm to assess ferrets’ ability to detect repetitions embedded in noise. Repeated embedded noise stimuli were composed of two overlapping continuous streams of brief (250- or 300-ms) broadband noise samples. The noise samples had second-order statistics (*i.e*., spectral and temporal envelope correlations) matched to natural sounds (23). Consistent with the goal of this study, the only streaming cue was repetition. These stimuli lacked other conventional streaming cues such as harmonicity and common onset time.

During the initial, *random phase* of each trial, samples for both streams were drawn randomly from a pool of 20 distinct noise samples (1-2.5-sec duration, Figure 1B). When all the samples are drawn randomly, they are perceived as a single stream. The random phase was followed immediately by the *repeating phase*, in which a target noise sample, different for each behavioral block, started to repeat in one sequence but not in the other. In humans, this repetition leads to perceptual separation of the two sequences into discrete streams (23). We refer to the sequence that contains the repeating target sample as the *foreground stream*, and the concurrent sequence with no repetition as the *background stream* (Figure 1B).

**Figure 1:**
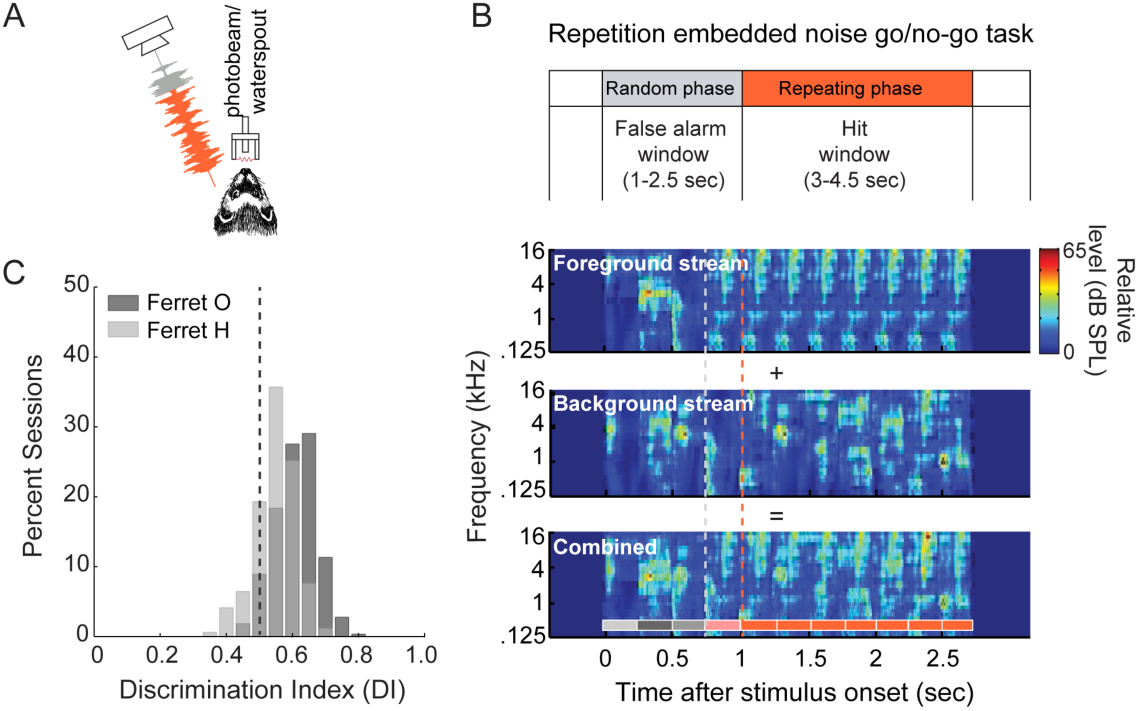
Ferrets are sensitive to repetitions embedded in mixtures. **A.** Ferrets were trained to respond to sound repetition by licking a waterspout. **B.** Schematic of the go/no-go task and spectrograms of repetition embedded noise stimuli from an example behavioral trial. Animals were exposed to the combination (bottom spectrogram) of two overlapping streams: a foreground stream containing a target sample (top), and a background stream, a non-repeating sequence of noise samples (middle). In this example, the target sample (orange boxes, bottom panel) starts repeating after three random noise samples (grey boxes). The grey dashed line marks the first occurrence of the target sample (pale orange), which is included in the random phase for analysis. The transition between random and repeating phase is marked by the orange dashed line and occurs when the target sample is first repeated. Animals were trained to withhold licking from a waterspout during the random phase (4-6 sec). To receive a water reward, they had to lick the waterspout following repetition onset. **C.** Distribution of discrimination index (DI) across behavior sessions for ferret O and ferret H after training was completed. Dashed line (0.5 DI), indicates chance performance.

Two ferrets (O and H) were trained to report when they detected the repetition of the target using a go/no-go detection paradigm. Head-fixed animals were required to withhold from licking a waterspout during the random phase and to lick after the onset of the repeating phase (Figure 1A-B). In each behavioral block (∼50-100 trials), two noise samples were chosen as targets from a pool of 20, each with 50% chance of occurring in a trial. Changing the identity of the targets between blocks avoided overtraining on a specific target. To measure behavioral performance in a task with continuous distractors and variable target times, we used a discrimination index (DI). This metric uses hit rate, false alarm rate, and reaction time to compute the area under the receiver operating characteristic (ROC) curve for target detection (29, 30). A DI greater than 0.5 indicates above-chance behavior. Both ferrets were able to learn the task and perform above chance within two months of training, indicating that they were able to perceive the repeating noise stream (Ferret O: mean DI = 0.61±0.004 SEM, *n* = 327; Ferret H: mean DI = 0.55±0.005 SEM, *n* = 171; Figure 1C).

### Neuronal responses are suppressed during the repeating phase

We recorded multi- and single-unit neural activity in primary (A1, *n* = 152) and secondary (PEG, *n* = 138) regions of the auditory cortex of five ferrets passively listening to the task stimuli. Two animals were trained on the repetition embedded noise task (behavior described above), and three were naïve to the task. Although all physiological data presented here were recorded in passively-listening ferrets, for consistency we refer to the same trial structure terminology as in the previous section. During electrophysiological recordings, one of the target noise samples was chosen to match each unit’s tuning (eliciting a relatively strong response) while the other was chosen at random. The two targets had equal probability of occurring on each trial.

To investigate the neurophysiological underpinnings of streaming due to repetition, we first looked at the raw firing rates of the recorded units in response to the repeated noise stimuli. Given the enhanced representation of repeating stimuli observed in behavioral experiments (22, 23, 31), we reasoned that evidence for the selective enhancement of foreground representation should be observed at the level of the auditory cortex. If this were true, we would expect the neural response to a target sample to change between random and repeating contexts.

To test this prediction, we computed the average peristimulus time histogram (PSTH) response across all occurrences of the target noise samples in the random phase (excluding any occurrences during first 250ms of the trial), and compared it to the average PSTH response to a balanced number of targets in the repeating phase (Figure 2A). Since the background sample was randomly selected for each presentation of the target, responses to the background sample were averaged out, and the PSTH primarily reflected responses to the target. To quantify changes in the response, we computed the gain term that scaled the PSTH for the random phase to best match the PSTH for the repeating phase. To allow for a direct comparison between gains generated by encoding models (see below), gain terms were log-transformed. Thus, negative values indicated suppressed responses during repetition and positive values indicated enhanced responses. For most units in A1 and PEG, log gain was less than zero (Figure 2B), indicating that the average target response in the repeating phase was suppressed with respect to the average target response in the random phase. Considering previous observations of neural adaptation to repeated stimuli in auditory cortex (24, 25), a decreased response to the target in the repeating phase is not unexpected.

**Figure 2:**
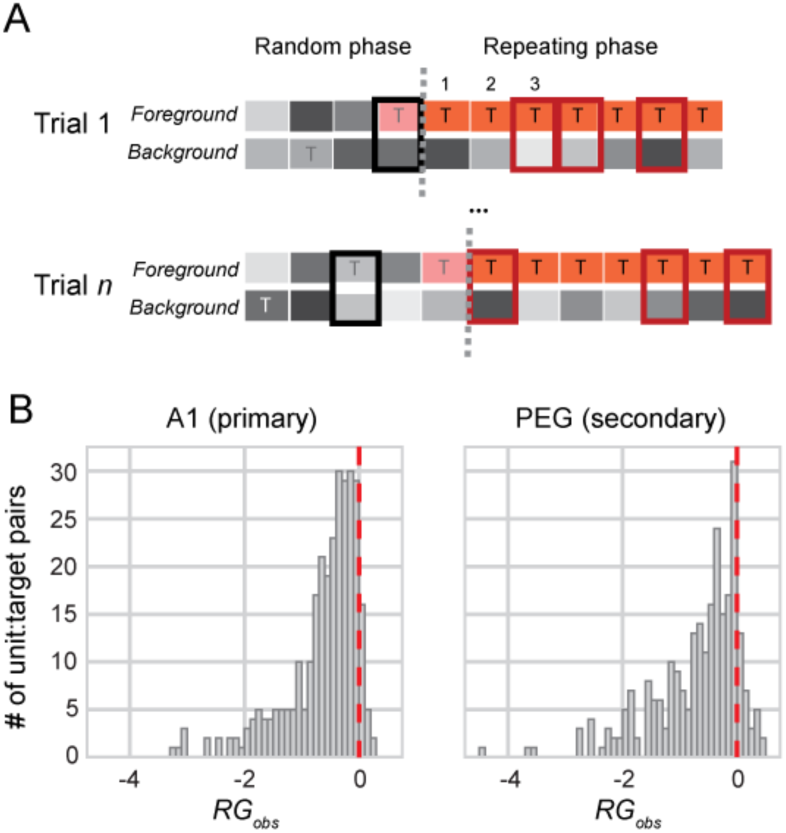
Activity in both A1 and PEG was suppressed during the repeating phase. **A.** Schematic of two trials. The background stream consisted entirely of random noise samples (gray). The foreground stream contained random samples during the random phase, which sometimes included the target sample (T). The final pair of noise samples in the random phase includes the target (light orange) as it had not yet begun to repeat. Average PSTH responses to each pair of samples that contained the target (thick rectangles) were computed separately for the random phase (black) and repeating phase (dark red). Pairs containing a target sample in the repeating phase were excluded to ensure that the number of appearances match between random/repeating (see *Material and Methods*). **B.** Distribution of observed repetition gain (*RG*_*obs*_) for A1 and PEG. The majority of target responses were suppressed (*RG* _*obs*_ < 0) during the repeating phase. Red dashed line indicates 0 (*i.e*., no difference between phases). Since results may depend on how well the unit responded to the target, all analyses of neural responses were performed separately for each unique unit-target pair (*n* = 304 A1, *n* = 276 PEG). Median *RG* _*obs*_ values, A1: −0.64 (95% CI [−0.74, −0.54]); PEG: -0.70 (95% CI [-0.81, 0.60]).

### Relative enhancement of responses to the repeating foreground stream

Simply comparing the average neural response to the target in the repeating phase to the random phase does not provide insight into any stream-specific effect that might emerge as a consequence of the repetition. To test for evidence of streaming in the neural response, we needed to independently assess the responses to the two streams. We reasoned that, even if the total response was suppressed, activity in the foreground stream in response to the repetition could be enhanced or suppressed *relative to* the background stream.

To test this prediction, we developed an encoding model in which the neural response was computed as the sum of responses to samples in each stream (*stream-dependent model*, see *Materials and Methods*). Using regression analysis, the relative contribution of each noise sample to the PSTH response was computed from the response to the noise streams. In the random phase the response was modeled as the sum of responses to each of the two concurrent samples and a baseline firing rate (*Eqn. 1*). In the repeating phase, the response to each sample was scaled according to whether it occurred in the foreground or background stream before summing (respectively, gain terms *RG*_*fg*_ and *RG*_*bg*_, *Eqn. 3*). We compared this model to a *stream-independent* model, in which responses to samples in both streams were scaled equally by a single gain term in the repeating phase (*RG*_*global*_, *Eqn. 2*). Responses were predicted relative to baseline firing rate, and the gain terms were log-scaled. Thus, positive gain indicates stronger modulation of the unit’s response (*i.e*., greater excitation and inhibition), and negative gain indicates weaker modulation of the unit’s response relative to baseline firing rate.

Figure 3 plots the average response to the target stimuli and predictions by the stream-dependent model for example units in A1 (top) and PEG (bottom). In both examples, the repetition gain for the background stream was negative (*RG*_*bg*_ = −1.0 in A1 and −1.6 in PEG). This means that neural responses to the background stream in the repeating phase were suppressed 64% and 80% relative to the random phase in A1 and PEG, respectively (Figure 3, black dotted line). Conversely, the foreground repetition gain term positively scaled the target sample response in the repeating phase (*RG*_*fg*_ = 0.47 in A1 and 0.79 in PEG; blue dashed line), increasing the response compared to the random phase by 60% and 79% increase in A1 and PEG, respectively (Figure 3, orange solid line).

**Figure 3:**
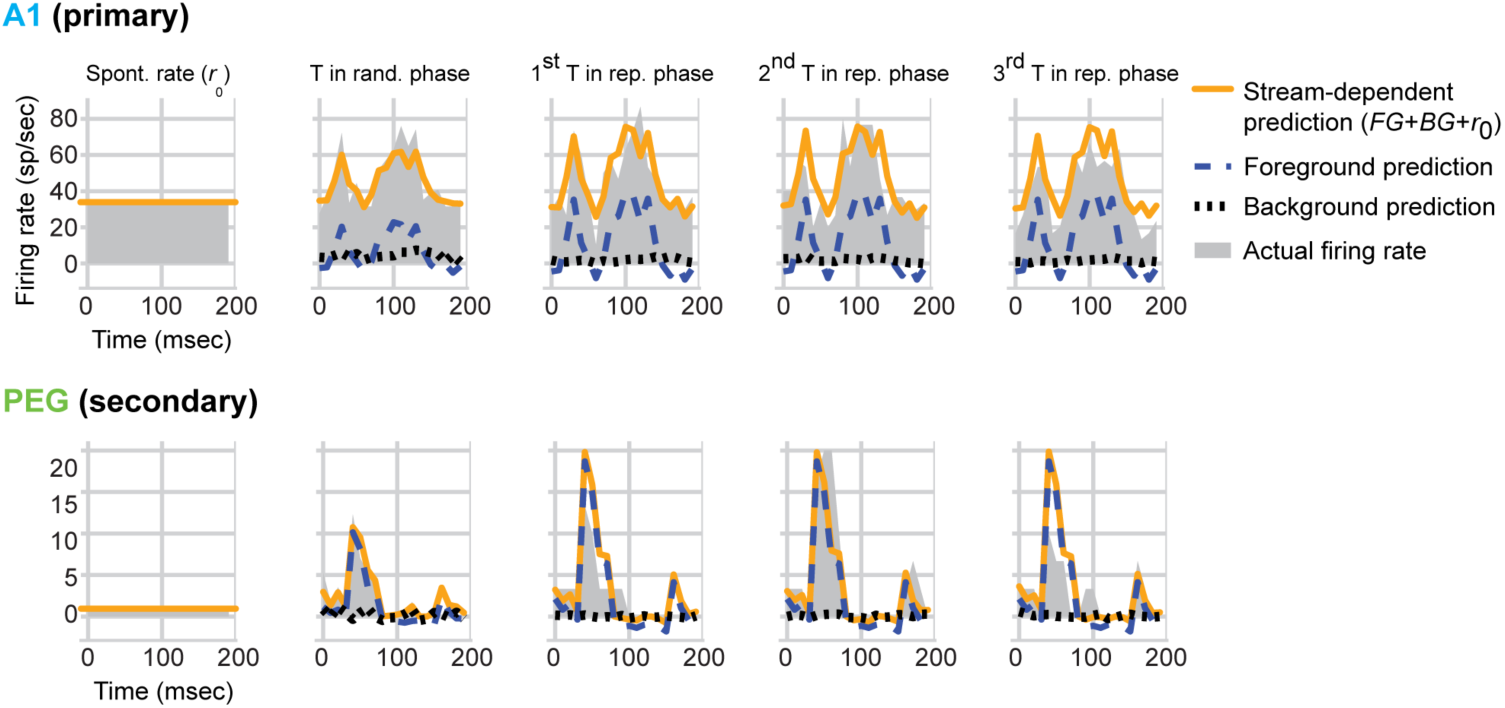
Example activity A1 and PEG. Example PSTH responses of units in A1 (top) and PEG (bottom) to the target sample (T) in the random versus repeating phase. Spontaneous rate (*SR*) is shown (1st column) for reference. Predictions from the stream-dependent model (orange) broken down into the contribution of the foreground (blue, dashed) and background (black, dotted) streams, shown for the average responses to the target in the random phase (2nd column), and the first three repetitions of the target sample in the repeating phase (3rd, 4th, and 5th columns).

Across the population, repetition gain was negative in the majority of unit-target pairs for both foreground and background streams (Figure 4A; A1: *n* = 304 unit-target pairs, mean *RG*_*bg*_ = −0.609 ± 0.041 SEM, mean *RG*_*fg*_ = −0.485 ± 0.035 SEM; PEG: *n* = 276, mean *RG*_*bg*_ = −0.935 ± 0.038 SEM, mean *RG*_*fg*_ = −0.518 ± 0.040 SEM). Similarly, in the stream-independent mode, units in both A1 and PEG usually had a negative *RG*_*global*_ (Figure 4B). This global suppression was consistent with the decrease observed in the average target response described above (Figure 2B; *r* = 0.63 between *RG*_*global*_ and target response gain, *p* < 0.0001, Student’s *t*-test).

**Figure 4:**
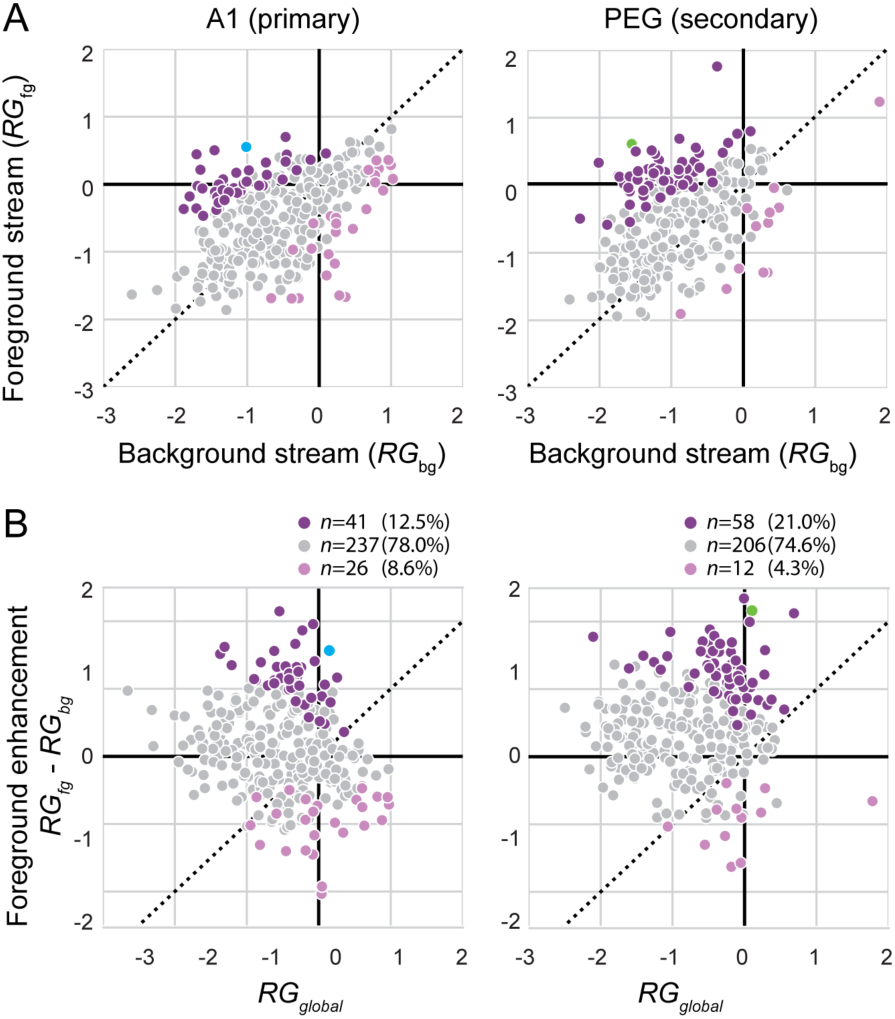
Selective foreground enhancement in A1 and PEG. **A.** Foreground (*RG*_*fg*_) versus background (*RG*_*bg*_) repetition gain measured in the stream-dependent model in A1 and PEG. Color indicates unit-target pairs in which values of *RG*_*fg*_ are significantly higher (dark purple) or lower (light purple) than values of *RG*_*bg*_ (95% credible interval for the difference does not overlap with 0). Grey indicates no significant difference. Dashed line indicates equality. **B.** Foreground enhancement (*RG*_*fg*_ − *RG*_*bg*_) plotted against overall gain change (*RG*_*global*_) during the target phase for A1. Colors are as in B. Number of data points with significant foreground enhancement, significant background enhancement and no change are shown above each plot. Mean foreground enhancement was significantly greater than zero in both areas. Data for examples in Figure 3 are highlighted for A1 (cyan) and PEG (green).

To test for relative enhancement of responses to the repeated foreground, we measured *foreground enhancement*, the difference between foreground and background repetition gain in each fit of the stream-dependent model (Figure 4B). Foreground enhancement was considered significant if the 95% confidence interval for the fitted parameter did not bracket 0 (see *Materials and Methods*). A subset of unit-target pairs displayed significant foreground enhancement (41/304 in A1, 58/276 in PEG, Figure 4B), meaning that in the repeating phase responses in the foreground stream were less suppressed or enhanced relative to responses in the background stream. In contrast, fewer units showed foreground suppression in either area (26/304 in A1, 12/276 in PEG). Across the set of unit-target pairs, mean foreground enhancement was significantly greater than zero in A1 (0.124, *p* = 0.004, Wilcoxon signed-rank) and PEG (0.416, *p* < 0.0001, Wilcoxon signed-rank) (Figure 4B). Mean foreground enhancement was stronger in PEG than in A1 (*p* < 0.0001, independent two-sample *t*-test).

Despite the overall suppression of activity during the repeating phase, these results support a model of selective enhancement of responses to the repeated foreground stream, consistent with the enhanced perception of the repeated stream relative to the random background (23).

### Auditory tuning properties predict the degree of repetition enhancement

Next, we wondered if the units showing significant foreground enhancement had distinct sensory encoding properties. For each unit, we quantified lifetime sparseness, a measure of selectivity for any one sample relative to the others (see *Materials and Methods, Eqn. 4*) (32). This metric is bounded between 0 and 1, where 0 indicates low sparseness (equal responses to all stimuli) and 1 indicates high sparseness (non-zero response to only one stimulus). Example responses to each noise sample in our collection for a unit with roughly average sparseness are plotted in Figure 5A. For each unit-target pair, we also computed target preference, the ratio of evoked response to the target sample versus the average response to all samples (see *Materials and Methods, Eqn. 5*). A target preference of 1 indicates that the modulation by the target is equivalent to the average response for all samples.

**Figure 5:**
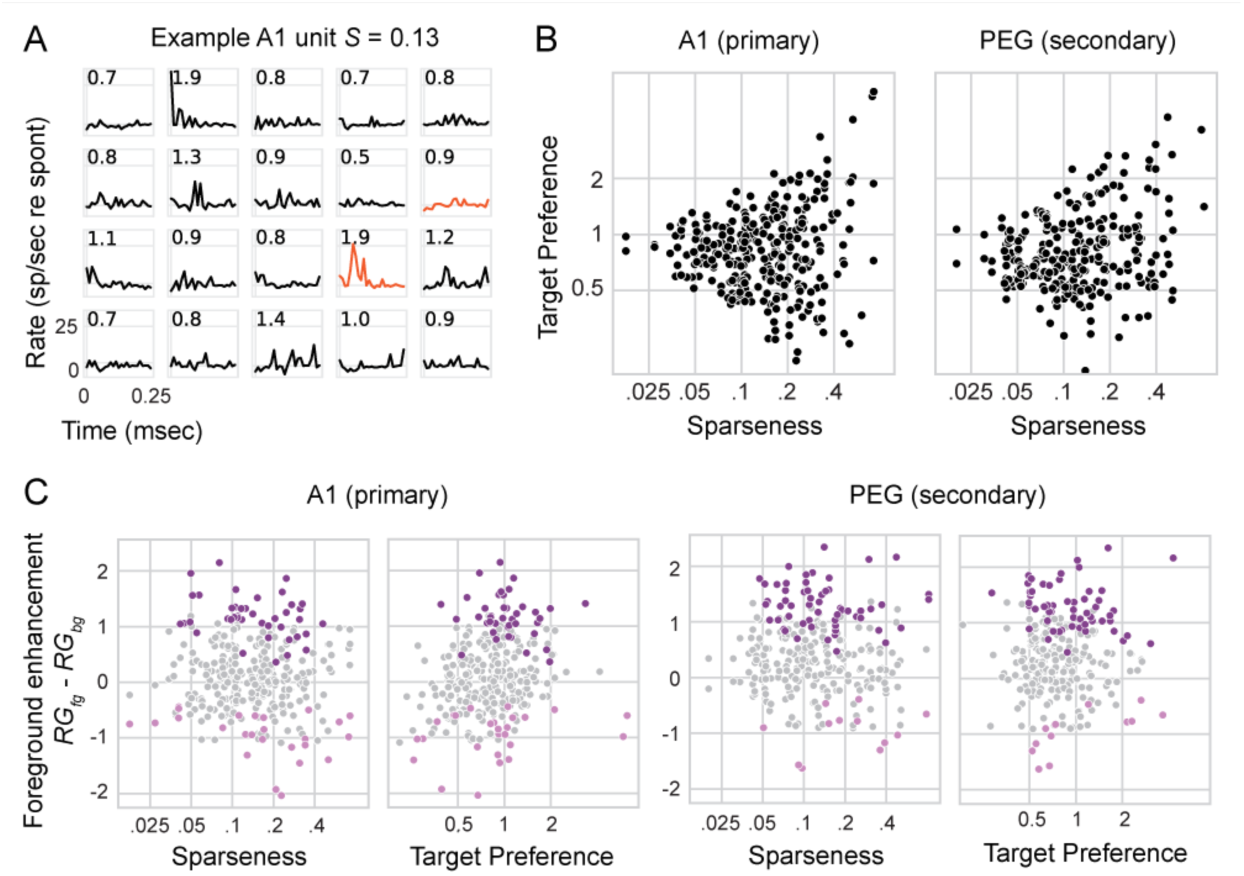
Relationship between target preference and sparseness in A1 and PEG units. PSTH responses to each of the 20 noise samples presented individually to a unit with relatively high sparseness (*S* = 0.13). Units such this one, responded well to only a few samples. Numbers indicate unit’s preference for that particular sample with respect to the others. Responses to target samples are indicated in orange. **B.** Scatter plot of target preference versus lifetime sparseness for each unit recorded from A1 and PEG. Target preference quantifies the response of a given unit to a target sample compared to the other 19 noise samples. Lifetime sparseness measures selectivity for the noise samples. Values of sparseness near 0 indicate units that responded similarly to all noise samples, and values near 1 indicate units that responded preferentially to a small number of samples. Units with high sparseness tended to have a greater variability in target preference. **C.** Scatter plots of foreground enhancement as a function of target preference and sparseness in A1 and PEG. Because sparseness and target preference are correlated, possible relationships with foreground enhancement were tested using a regression model, which is detailed in Table 1. No significant relationship was observed between sparseness and foreground enhancement in either area (*p* > 0.05, linear hypothesis test (75)). However, units in A1 with strong target preference tended to show stronger foreground enhancement (A1: *p* < 0.0001, PEG: *p* = 0.06; linear hypothesis test).

The relationship between each unit’s sparseness, target preference, and auditory area (A1 or PEG) and its foreground enhancement (Figure 5C) was quantified by a general linear mixed model (see *Materials and Methods, Eqn. 6*) with area, target preference, and sparseness as fixed effects and unit as a random effect. All two- and three-way interactions between the fixed parameters were included, and complete results are shown in Table 1. This model identified a significant relationship between target preference and foreground enhancement in A1 (*e.g*., for every increase in target preference by 1, foreground enhancement increased by 0.46), which was significantly modulated by sparseness. That is, foreground enhancement was stronger in units with high target preference, and this effect decreased with increasing sparseness (Figure 5C; *e.g*., for a unit with a sparseness of 0.05, the effect of target preference on foreground enhancement would be 0.41, whereas for a unit with a sparseness of 0.4, the effect of target preference on foreground enhancement would be 0.04). In contrast, in PEG there was no significant relationship between foreground enhancement and either sparseness, target preference, or the interaction of target preference and sparseness (Table 1).

**Table 1:** Regression analysis of auditory tuning effects on foreground enhancement. Results of the mixed linear model for foreground enhancement, with target preference and sparseness as fixed effects, computed separately for A1 and PEG data. Significance was assessed using a two-tailed *t*-test.

| area | effect | coef. | coef | std err | z | P>\|z\| | Conf. Int. Low | Conf. Int. Upp. |
| --- | --- | --- | --- | --- | --- | --- | --- | --- |
| A1 | intercept | $\beta_0$ | 0.15 | 0.04 | 3.57 | 0.0004 | 0.07 | 0.23 |
| | sparseness | $\beta_2$ | -0.42 | 0.32 | -1.33 | 0.1835 | -1.04 | 0.20 |
| | target preference | $\beta_4$ | 0.46 | 0.11 | 4.38 | 0.0000 | 0.26 | 0.67 |
| | target preference * sparseness | $\beta_4$ | -1.04 | 0.27 | -3.92 | 0.0001 | -1.57 | -0.52 |
| PEG | intercept | $\beta_0 + \beta_1$ | 0.42 | 0.04 | 9.42 | 0.0000 | 0.33 | 0.50 |
| | sparseness | $\beta_2 + \beta_3$ | -0.32 | 0.32 | -0.98 | 0.3293 | -0.95 | 0.32 |
| | target preference | $\beta_4 + \beta_5$ | 0.05 | 0.10 | 0.50 | 0.6167 | -0.14 | 0.24 |
| | target preference * sparseness | $\beta_4 + \beta_7$ | -0.11 | 0.35 | -0.30 | 0.7644 | -0.80 | 0.59 |

Thus, in A1, units that responded to many stimuli (low sparseness) but had a relatively strong preference to a target (high target preference) tended to show the most foreground enhancement. In PEG, enhancement was stronger overall and affects responses more uniformly, regardless of auditory selectivity. These differences between PEG than A1 suggest a gradual emergence of repetition-related streaming along the cortical auditory pathway.

### Foreground enhancement increases accuracy of spectro-temporal receptive field models

To validate the gain changes observed in the PSTH-based model and to quantify their effect on sound-evoked activity, we modeled the same data with a spectro-temporal receptive field (STRF). In the classic linear-nonlinear (LN) STRF (see *Material and Methods, Eqns. 7-9*), the time-varying neural response is modeled as a linear weighted sum of the stimulus spectrogram (33, 34). We developed a context-dependent model, in which spectrograms for each stream were scaled separately by a gain term before input to the STRF (*Eqns. 10-11*). This stream-dependent scaling followed the same logic as the PSTH-based model described above. That is, a separate spectrogram for each stream was scaled by free parameters that depended on phase (random or repeating) and, during the repeating phase, stream identity (foreground or background). The rescaled spectrograms were summed and provided input to a traditional LN STRF. Context gain parameters and STRF parameters were fit simultaneously (Figure 6A) (28). We refer to this model as the *phase*+*stream* STRF.

**Figure 6:**
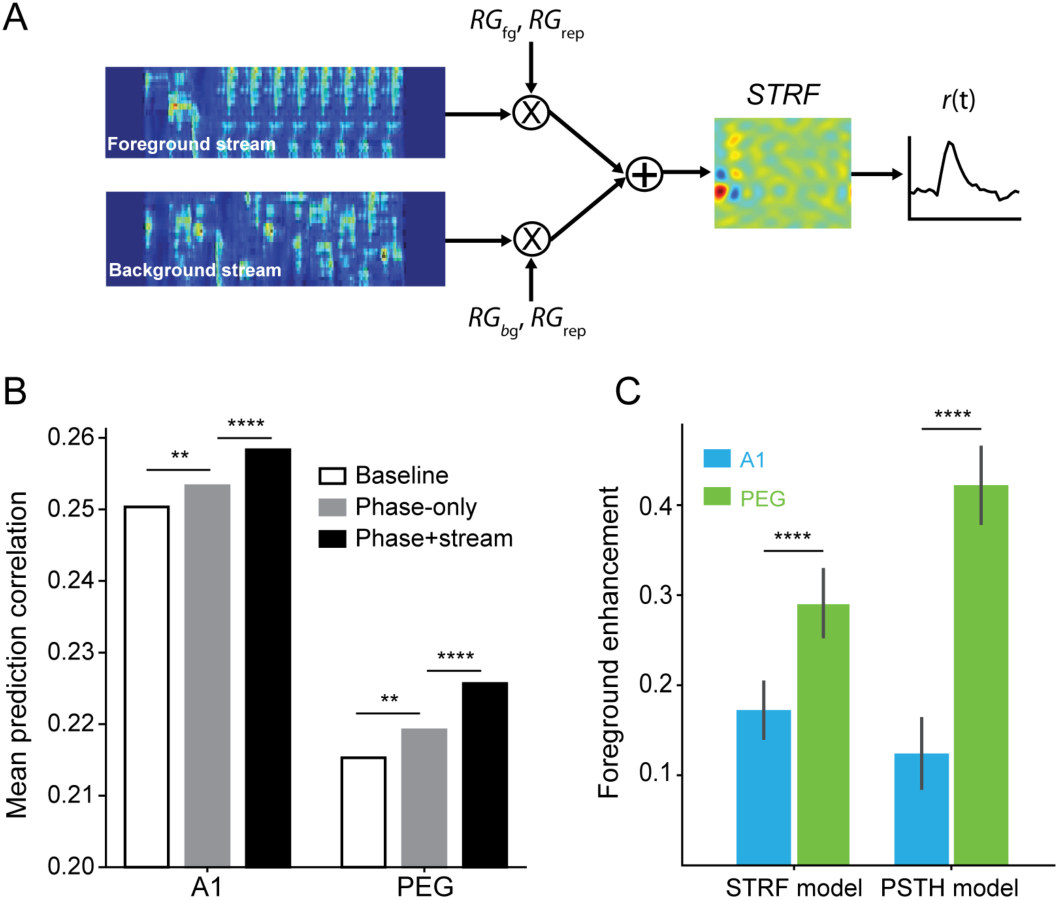
STRF-based model corroborates PSTH-based model findings of stream-specific gain changes in A1 and PEG. **A.** Schematic of the spectro-temporal receptive field (STRF)-based encoding model. Spectrograms of the foreground and background streams are scaled by context-dependent gain. Separate gain terms are applied to the foreground and background streams during the repeating phase. The sum of the scaled spectrograms provides input to a traditional linear-nonlinear STRF, which predicts the time-varying response to the repeated noise stimuli, *r*(*t*), as a weighted sum of the spectrogram. **B.** Mean prediction correlation coefficient (Pearson’s *r*) between the predicted and actual time-varying neural response for A1 and PEG units, plotted for the *baseline* STRF-based model (both stream identity and repetition permuted in time prior to fitting), for the *phase-only* model (stream identity shuffled), and for the full *phase*+*stream* model. Asterisks indicate *p*-values associated with Wilcoxon signed-rank test within-area model comparisons (A1, *n* = 152; PEG, *n* = 138). **C.** Mean foreground enhancement for A1 and PEG and for STRF- and PSTH-based models. Asterisks indicate *p*-values associated with independent two-sample *t*-test between values of foreground enhancement in A1 and PEG. \*\**p* <0.01, \*\*\*\**p*<0.0001.

We used 20-fold cross validation to compare the prediction accuracy of the phase+stream STRF to two control models: a *phase-only* STRF, in which stream identity was shuffled in time before fitting, and a *baseline* STRF, in which both phase and stream identity were shuffled. The phase-only STRF accounted for changes in gain due to repetition but independent of foreground versus background stream identity, analogous to the stream-independent model above. The phase+stream STRF predicted time-varying responses more accurately than the phase-only STRF in both A1 and PEG, confirming a significant influence of stream identity on relative gain (A1: *p* < 0.0001, PEG: *p* < 0.0001, Wilcoxon signed-rank test, Figure 6B).

To measure the relative enhancement of the two streams, we compared the stream-specific gain terms from the model fits, equivalent to *RG*_*fg*_ and *RG*_*bg*_ discussed above. We observed a significant relative increase in foreground versus background gain in both A1 (mean foreground enhancement 0.159; *p* < 0.0001, Wilcoxon signed-rank test) and PEG (mean 0.291; *p* < 0.0001, Wilcoxon signed-rank test). This result provided further evidence for stream-dependent changes in gain. These changes in gain followed the same pattern as mean foreground enhancement in the stream-dependent model above (correlation coefficient measured between foreground enhancement for PSTH- and STRF-based models in A1: Pearson’s *r* 0.39, *p* < 0.0001 and PEG: Pearson’s *r* 0.22, *p* = 0.0005; Figure 6C).

The comparison of phase-only and baseline STRFs measured the effect of repetition alone on evoked activity (independent of stream identity). On average, the phase-only STRF had greater prediction accuracy than the baseline STRF in both areas (A1: p < 0.0001, PEG: p < 0.0001, Wilcoxon signed-rank test, Figure 6B). Thus, the STRF-based models confirmed an effect of repetition on evoked activity. Moreover, overall gain was suppressed during the repeating phase (mean A1: −0.11, PEG: - 0.042, data not shown), as observed in the PSTH-based models above (Figure 4). Thus, this approach provides additional evidence for a streaming mechanism in which repetition leads to overall suppression of the neural responses, but with less prominent suppression of the foreground stream relative to the background.

## Discussion

In natural environments, temporally co-varying sound features tend to be grouped by the brain into a single object (35). Sound repetition is one such feature that can induce stream segregation in human listeners. Subjects are able to identify individual, previously unheard noise samples if they are repeated simultaneously to a mixture of different non-repeating samples (23). The goal of the current study was to investigate the neural underpinnings of streaming cued by sound repetition. We developed an animal model for repetition detection and found evidence for enhanced representation of a repeating foreground stream in single-unit activity in auditory cortex. This representation appears to emerge hierarchically, as streaming effects were stronger in secondary (PEG) than in primary (A1) auditory cortical fields.

### Mechanisms of repetition-induced stream segregation

Previous studies that have explored the neural signature of streaming at the single-unit level have primarily used alternating sequences of pure tones (36, 37). Relevant to the current study, Micheyl and collaborators presented sequences of “ABA_” tone triplets to awake macaques and examined the pattern of activity evoked in A1 (36). Tone A was chosen to be on the best frequency of the recorded unit, while tone B was placed at a frequency of 1-9 semitones from tone A. The authors found that, even if responses to both tones decreased relative to their presentation in isolation, responses to the non-preferred B tones decreased to a greater extent. In the current study, we observed a similar effect, that relative enhancement of the foreground stream was more pronounced in units well-tuned to the repeated noise sample. Thus, our results are consistent with observations based on the ABA tone paradigm. They also provide evidence that the same principles generalize to streaming of simultaneous, naturalistic sounds, a situation that more closely relates to animals’ everyday sound experience.

Sound features that belong to the same source tend to begin and end at the same time. This phenomenon has been formalized for streaming in the temporal coherence model (17, 18). Teki *et al.* (2016) demonstrated that human listeners are highly sensitive to repetition of sounds presented in the context of a random mixture of chords. Similar to our findings, the authors observed that repeating sounds tend to fuse together into a “foreground” that emerges from a randomly changing background (38, 39). Here, we propose that foreground enhancement contributes to streaming repeating sounds in the context of a random background.

However, it is important to note that an expectation of enhanced responses to “foreground” stimuli may reflect a biased expectation. There is no *a priori* requirement for sounds that perceptually pop out as a foreground to evoke an enhanced (or less suppressed) neural response. For example, Bar-Yosef and collaborators, investigated the interactions in A1 of anesthetized cats during simultaneous presentation of bird chirps and background noise, simulating a naturalistic auditory scene (40, 41). To their surprise, neural responses were in fact dominated by the background noise, despite it being presented at a lower intensity than the foreground bird chirp. The authors interpreted this finding in an evolutionary context, in which it is advantageous for prey to pay attention to subtle changes in the background to avoid predators which might be using foreground sounds to mask the sound of their own approach. This example suggests that the brain might enhance different components of a sound depending on the context and identity of that sound. Thus, it is important to consider the traditional ecosystem niche of the animal model when interpreting our findings. More experiments testing streaming effects in natural contexts will be needed to further elucidate how the brain streams repeated sound features with behavioral relevance.

### Streaming analysis

To capture differences in responses to simultaneous repeated and non-repeated noise samples, we relied on model predictions. In this paradigm, the neural response is the sum of responses to two simultaneous stimuli, and the component responses cannot be separated in the raw neural firing rate. Therefore, we constructed encoding models that teased apart stream-dependent activity computationally. This analysis showed that, even though most neural responses were suppressed by repetition—likely due to adaptation (24, 42, 43), responses to the foreground stream where less suppressed than the background or even enhanced. This approach established a methodology that could be used in other datasets where there is a need to separate effects on neural responses to simultaneously occurring inputs.

The challenge of separating responses to simultaneously occurring sounds has been previously addressed for neural population activity using a similar modeling approach (44). Ding and Simon asked human subjects to listen to one of two competing speakers and recorded brain activity via magnetoencephalography (MEG). To investigate the neural encoding process, they fit a separate STRF (or more precisely a “TRF”, since the MEG data did not resolve spectral tuning) for each of the two simultaneously presented speech streams. Neural activity was found to preferentially synchronize to the speech envelope of the attended speaker. Furthermore, the latency and source location of the two components suggested a hierarchy of auditory processing in which the representation of the attended object emerges from core to posterior auditory cortex (44). These results are largely consistent with the foreground enhancement observed in the current study, suggesting that top-down attention and bottom-up pop-out effects could be mediated by common mechanisms.

Another approach used to investigate the neural signature of streaming is by stimulus decoding, or reconstruction (12, 44, 45). A decoding model describes the relationship between stimulus and response similarly to the STRF, but in the opposite direction. That is, decoding uses neural population activity to reconstruct the sound stimulus input. If the reconstruction of the envelope has a higher correlation to the envelope of the attended stream rather than the non-attended stream or the two streams combined, it would suggest enhanced coding of the attended stream. In a human MEG study, Ding and Simon (2012) found that this was indeed the case. Similar results were also obtained by Mesgarani and Chang (2012) using data collected from non-primary auditory cortex via electrocorticography (ECoG) (12).

In our study, a decoding analysis could complement the encoding approach, potentially revealing how the relative enhancement of the repeating stream allows for the separation of the two streams. Specifically, we would predict that for units with positive foreground enhancement stimulus reconstruction would be more accurate for the foreground stream compared to the background stream, matching perception.

### Relation of repetition enhancement to stimulus-specific adaptation

The ability of the brain to detect regularities is not only crucial for identifying an auditory object embedded in a noisy scene, but also for making predictions about the environment, thereby making the system sensitive to deviance (6, 19). Substantial effort has been devoted to understanding the mechanisms of deviance detection, with a focus on the mismatch negativity (MMN) in human studies (27) and on stimulus-specific adaptation (SSA) in animals (24, 25, 46). Evidence for SSA has been found in the inferior colliculus and thalamus (47–49), but the first lemniscal region in which SSA has been shown to be prominent is A1 (50, 51). Mechanistically, SSA is thought to arise from a combination of feedforward synaptic depression and local cortical inhibition (52–54).

Selective enhancement of the neural response to a repeating sound might seem like an intuitive prediction, based on behavioral studies of repetition-induced streaming (22, 23). However, this enhancement may be surprising when viewed in the context of SSA (25, 55). If SSA affects responses to simultaneous stimuli the same way as responses to sequential stimuli, one would expect a relative suppression of responses to the foreground stream in repeated embedded noise. However, our results show the opposite effect, *i.e*., relative suppression of the non-repeating background stream, especially for target-preferring neurons. We propose that while SSA can account for the overall decreased response to both streams (24, 43), a separate mechanism must be responsible for the further suppression of sounds that occur simultaneously to the repeating foreground. Furthermore, the fact that foreground enhancement is more prominent in secondary auditory cortical fields (PEG) than primary areas (A1), suggests a hierarchical mechanism by which the enhancement emerges along the auditory cortical pathways.

### Animal models for streaming

Most behavioral studies of auditory streaming have been performed in humans (2–4, 56, 57). This is not surprising, as perceptual measurement of streaming in nonhuman species is challenging (for recent reviews on nonhuman behavioral studies of auditory streaming see 58, 59). Within a small number of such animal studies, however, the ferret has been identified as a model for streaming of alternating tone sequences and tone clouds (16, 60), and had been used to study its neurophysiological bases (18).

Here, we developed the ferret as a model for streaming repeated sequences of simultaneously presented complex sounds. We designed an auditory task where animals had to report the occurrence of a repetition emerging from random overlapping noise samples. Ferrets were able to perform this task, suggesting that they could perceive repetition of complex sound features as a distinct component of the stimulus. Since the identity of the repeated sample was changed across behavioral blocks, we could exclude the possibility that the animals used specific spectro-temporal features of the target sample to perform the task. While the ability to report the occurrence of repetitions in one stream is not a direct proof that ferrets perceived two separate streams in the same way as humans, it confirms that they did perceive the occurrence of repetitions.

### The role of attention in repetition-induced streaming

Our physiological experiments were conducted on passively listening ferrets without explicit control of attention. While attention is known to modulate sensory responses across multiple brain areas (61–63), the role of attention on repetition-based streaming is controversial. Masutomi, McDermott *et al*. tested this question directly by asking human subjects to perform the same task as in McDermott *et al*., 2011, but while also performing a decoy visual task (31). The authors found that human listeners were equally able to recover the identity of the repeating noise sample even when their attention was directed away from the sound, indicating that repetition-based streaming is a bottom-up process.

Several other studies have shown that human listeners are extremely sensitive to regular patterns rapidly emerging from complex sequences of sound (38, 64). Barascud *et al*. investigated how human listeners discover temporal patterns and statistical regularities in complex sound sequences (64). They found that subjects’ behavior matched the one of an ideal observer, even when distracted by a decoy visual task, again suggesting that detection of sound repetition might be a phenomenon that does not require attentional focus.

Streaming of more complex sounds (e.g., speech) is facilitated by directing attention to specific sound features that distinguish a foreground from a background (65). For example, Mesgarani & Chang, 2012 presented human listeners with two streams of speech. What was referred to as foreground or background changed across trials in response to a specific word that cued participants to either listen to the female or the male voice. The authors found that listeners were much better at reporting the content of sentences that they were cued to pay attention to with respect to non-cued sentences presented simultaneously. Furthermore, the signature of this “foreground enhancement” is present at the level of neural activity measured by ECoG (12) and MEG (44). Future experiments incorporating behavior into neurophysiological recordings may explain whether the pre-attentive foreground enhancement effects reported here are mediated by the same mechanisms as those that enhance actively attended streams.

## Materials and Methods

All procedures were approved by the Oregon Health and Science University Institutional Animal Care and Use Committee and conform to the United States Department of Agriculture standards.

### Surgical procedure

Animal care and procedures were similar to those described previously for neurophysiological recordings from awake ferrets (66). Five spayed, de-scented young adult ferrets (two females, three males) were obtained from an animal supplier (Marshall Farms, New York). Normal auditory thresholds were confirmed by measuring auditory brainstem responses. A sterile surgery was then performed under isoflurane anesthesia to mount a post for subsequent head fixation and to expose a 10-mm^2^ portion of the skull over the auditory cortex where the craniotomy would be subsequently opened. A light-cured composite (Charisma, Heraeus Kulzer) anchored a custom stainless-steel head-post on the midline in the posterior region of the skull. The stability of the implant was also supported by 8-10 stainless self-tapping set screws mounted in the skull (Synthes). The whole implant was then built up to its final shape with layers of Charisma and acrylic pink cement (AM Systems).

During the first week post-surgery, the animal was treated prophylactically with broad-spectrum antibiotics (10 mg/kg Baytril). For the first two weeks the wound was cleaned with antiseptics (Betadine and Chlorexidine) and bandaged daily. After the wound margin was healed, cleaning and bandaging occurred every 2-3 days through the life of the animal. This method revealed to be effective in minimizing infections of the wound margin.

### Stimuli and Acoustics

Repeated embedded noise stimuli used in the present study were generated using the algorithm from McDermott *et al*. (2011) (23). Brief, 250- or 300-ms duration samples of broadband Gaussian noise were filtered to have spectro-temporal correlations matched to natural sounds but without common grouping cues, such as harmonic regularities and common onsets (23). The spectral range of the noise (125-16,000 Hz or 250-20,000 Hz) was chosen to span the tuning of the current recording site. An experimental trial consisted of continuous sequences of 10-12 noise samples (0 ms inter-sample interval) drawn randomly from a pool of twenty distinct samples (Figure 1). The order of samples varied between trials. Either one stream of samples was presented (single stream trial) or two streams were overlaid and presented simultaneously (dual stream trial). At a random time (after 3-11 samples, median 6 samples), the sample in one stream (target sample) began to repeat. In dual stream trials, this repetition occurred only in one of the two streams, while samples in the other stream continued to be drawn randomly. In human studies, the repeating sample has been shown to pop out perceptually as a salient stream (23). Thus, the stream containing the repeated sample is referred to here as the *foreground*, and the non-repeating stream as the *background* (Figure 1B). The period of the trial containing only random samples is referred to as the *random phase*, and the segment starting with the first repetition of the target sample, where the two streams perceptually diverge, is referred to as the *repeating phase* (Figure 1B). With the exception of the spectro-temporal receptive field analysis, the first sample of the random phase was excluded from analysis to minimize the effect of onset-related adaptation on our analysis.

All behavioral and physiological experiments were conducted inside a custom double-walled sound-isolating chamber with inside dimensions of 8’ × 8’ × 6’ (L × W × H). A custom second wall was added to a single-walled factory chamber (Professional Model, Gretch-Ken Inc.) with a wooden frame and an inner wall composed of 3/4” MDF board. The air gap between the outer and inner walls was 1.5”. The inside wall was lined with 3” sound absorbing foam (Pinta Acoustics). The chamber attenuated sounds above 2 kHz by more than 60dB. Sounds from 0.2-2 kHz were attenuated 30-60 dB, falling off approximately linearly with log-frequency.

Stimulus presentation and behavioral control were provided by custom MATLAB software (Mathworks Inc.). Digitally generated sounds were D/A converted (100 kHz, National Instruments PCI-6229), and presented through a sound transducer (Manger W05) driven with a power amplifier (Crown D-75A). The speaker was placed one meter from the animal’s head, 30° contralateral to the cortical hemisphere under study. Sound level was calibrated using a 1/2” microphone (Bruel & Kjaer 4191). Stimuli were presented with 10ms cos^2^ onset and offset ramps.

### Behavior

Two ferrets (one male, ferret H, and a female, ferret O) were trained to report the occurrence of repeated target noise samples in the repeated embedded noise stimuli using a go/no-go paradigm (29). Starting two weeks after the implant surgery, each ferret was gradually habituated to head fixation by a custom stereotaxic apparatus in a plexiglass tube. Habituation sessions initially lasted for 5 minutes and increased by increments of 5-10 minutes until the ferret lay comfortably for at least one hour. At this time the ferret was placed on a restricted water schedule and began behavioral training. During training and physiological recording sessions that involved behavior, the ferret was kept in water restriction for five days/week, and almost all the daily water intake (40-80 ml) was delivered through behavior. Their diet was supplemented with 20 ml/day of high protein Ensure (Abbott). Water restriction was to be discontinued if weight dropped below 20% of the initial weight, but this did not happen with either ferret. Water rewards were delivered through a spout positioned close to the ferret’s nose. Delivery was controlled electronically with a solenoid valve. Each time the ferret licked the waterspout, it caused a beam formed by an infrared LED and photo-diode placed across the spout to be discontinued (Figure 1A). This system allowed us to precisely record the timing of each lick relative to stimulus presentation.

After trial onset, animals were required to refrain from licking until the onset of the repeating phase, *i.e*., after the occurrence of a repeated sample. Licks during the random phase were recorded as false alarms and punished with a 4-6 sec time-out. Licks that occurred in the repeating phase were recorded as hits and always rewarded with one to two drops of water (Figure 2.1B). Each behavioral session had two target samples whose identity varied from session to session to avoid ferret overexposure to a given target spectro-temporal features.

To shape the animal’s behavior, training started with a high signal-to-noise ratio (SNR) between random and repeating phases. SNR was then slowly decreased until 0dB SNR was reached. Parameters such as spectral modulation depth of the two streams and length of the random phase/false alarm window were also adjusted over the training period. Performance was assessed by a discrimination index (DI) computed from the area under the receiver operating characteristic (ROC) curve for detection of the target in the repeating phase (29, 30). DI combines information about hit rate, false alarm rate, and reaction time, and has a higher value for the higher, lower, and faster these scores are, respectively. A DI greater than 0.5 indicates above-chance performance. Criterion was reached as the ferret performed at DI > 0.5, with 0 SNR and 0 modulation depth difference for four consecutive days.

### Electrophysiology

Single- and multi-unit neural recordings were performed in the two trained animals and in three additional task-naïve animals. A small (∼1-2 mm diameter) craniotomy was opened over the auditory cortex, in a location chosen based on stereotaxic coordinates and superficial landmarks on the skull marked during surgery. Initial recordings targeted primary regions of the ferret auditory cortex (A1), and recording location was confirmed by characteristic short-latency responses to tone stimuli and by tonotopic organization of frequency selectivity (67). Recordings in secondary auditory cortex (PEG) were then performed in the field ventral to A1, identified by a reversal in the tonotopic gradient.

On each recording day, 1 to 4 high-impedance tungsten microelectrodes (FHC or A-M Systems, impedance 1-5 MΩ) were slowly advanced into cortex with independent motorized microdrives (Alpha-Omega). The electrodes were positioned (Kopf Instruments) such that the angle was roughly normal to the surface of the brain (∼28-40°). Stimulus presentation and electrode advancement were controlled from outside the sound booth, and animals were monitored through a video camera. Neural signals were recorded using open-source data acquisition software (MANTA, (68)) Raw traces were bandpass-filtered (0.3-10 kHz), amplified (10k, A-M Systems 1800 or 3600 AC amplifier), digitized (20 kHz, National Instruments PCI-6052E) and stored for subsequent offline analysis. Putative spikes were extracted from the continuous signal by collecting all events ≥4 standard deviations from zero. Different spike waves were separated from each other and from noise using principle component analysis and *k*-means clustering (69). Single units (>95% isolation) and stable multiunits (>70% isolation) were included in this study, resulting in a total of 141 A1 and 136 PEG units.

Between recording sessions, the exposed recording chamber surrounding the craniotomy was covered with polysiloxane impression material (GC America). After several electrophysiological penetrations (usually about 5-10), the craniotomy was expanded or a new craniotomy was opened to expose new regions of auditory cortex. When possible, old craniotomies were covered with a layer of bone wax and allowed to heal. Multiple craniotomies were performed on both hemispheres.

### Analysis

#### Effect of repetition on target responses

To assess the effect of repetition on overall responsiveness, we first measured changes in the response to the target sample between random and repeating trial phases. We computed the peristimulus time histogram (PSTH) response to each occurrence of target sample in the stimulus separately for the random phase and repeating phase, using data from dual-stream trials only. Spontaneous rate was subtracted from the PSTH to ensure the fraction term reflected changes in the evoked response. We then computed the gain term that minimized the least squares difference between evoked responses in the two phases. Log of the measured gain is reported to allow for direct comparison with the results of subsequent modeling analysis (see below).

#### PSTH-based models

Auditory cortical neurons could support segregation of the repeated stream either by changing the overall gain of their response to the repeating stream (stream-independent) or by differentially enhancing responses to one or the other stream (stream-dependent). To test these alternative predictions, we fit the data using *stream-independent* and *stream-dependent* models. In both models, responses were predicted using a weighted sum of time-varying responses to each noise sample. During the random phase, the time-varying response was the linear sum of a response to the foreground stream, response to the background stream, and spontaneous spike rate:

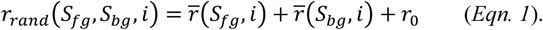

Here, *S*_*fg*_ and *S*_*bg*_ are the identity of samples in the foreground and background streams, respectively, and *r*_*0*_ is the spontaneous rate. 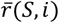 is the contribution of sample S to the evoked spike count in *i*-th time bin following sample onset.

For the stream-independent model, responses during the repeated phase were computed,

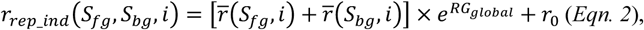

where *RG*_*global*_ scales responses to both streams. For the stream-dependent model, responses during the repeated phase were computed,

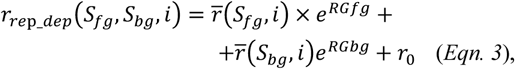

where *RG*_*fg*_ and *RG*_*bg*_ modulate the respective stream responses separately before they are summed. The use of an exponent simplifies interpretation of gain changes such that values of *RG* > 0 indicate enhancement and values of *RG* < 0 indicate suppression.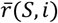 can be negative, which allows for suppressed responses relative to the spontaneous rate. In this case, if a unit has both enhanced and suppressed responses, *RG* will scale both responses equally (*e.g*., if *RG* > 0, there will be a decrease in spike rate during negative responses and an increase in spike rate during enhanced responses). The difference *RG*_*fg*_ − *RG*_*bg*_ is the relative enhancement between streams, here referred to as *foreground enhancement*. If *RG*_*fg*_ > *RG*_*bg*_, then the neural response to the foreground stream is enhanced relative to the background stream.

The repeating phase had many more presentations of the target sample than the random phase. To minimize potential bias when fitting the data, we randomly discarded target samples from the repeating phase such that the number of target samples in the repeating phase matched the number of target samples in the random phase. As mentioned earlier, the first sample of the random phase (*i.e.*, the very first sample in the trial) was excluded from analysis to minimize the effect of onset-adaptation in our analysis.

Models were fit to maximize Poisson likelihood of free parameters using Bayesian regression. A normal prior with mean 0 and standard deviation 10 was set on both *r*. and 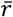. A normal prior with mean 0 and standard deviation 1 was set on all RG parameters. The model was fit three times using a different set of random starting values for each coefficient. Two thousand samples for each fit were acquired with a No-U-Turn Sampler, an extension to Hamiltonian Monte Carlo that eliminates the need to set a number of steps (70). Gelman-Rubin statistics were computed for each fit to ensure all the fits converged to the same final estimate 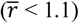.

The posteriors for *RG*_*global*_, *RG*_*fg*_ and *RG*_*bg*_ were extracted from the Bayes model. An *RG* parameter for which the 95% credible interval (as derived from the posterior) was less than 0 were considered to have significant suppression. Parameters with an interval greater than 0 were considered to have significant enhancement. For all data points shown, the means of the relevant posterior are plotted.

#### Lifetime sparseness and target preference

We quantified sparseness (*S*), a measure of unit selectivity for a given sample relative to the others in the collection (adapted from (32)),

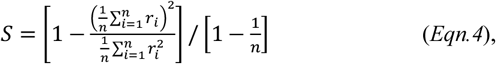

where *r*_*i*_ is the standard deviation of the PSTH (computed using the average of response to the token in the random phase of single-stream trials) for the *i*^th^ sample and *n* is the total number of noise samples. We quantified target preference (*TP*), a measure of how well the target sample modulates the unit’s response,

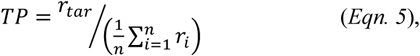

where *r*_*tar*_ is the standard deviation of the target PSTH and the other terms are defined as for sparseness. The use of standard deviation to measure response magnitude means that strong suppression or enhancement yield similar response strength.

To assess whether there was a significant effect of sparseness and/or target preference on foreground enhancement, we used a mixed linear model sparseness (S) and target preference (T) according to the following model design:

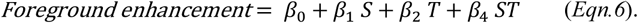

#### Spectro-temporal receptive field models

In addition to the PSTH-based models, which fit responses to individual noise samples, we confirmed that the same streaming effects were captured by a context-dependent spectro-temporal receptive field (STRF) model (28). The classic linear-nonlinear (LN) STRF models neural activity as the linear weighted sum of the preceding stimulus spectrogram, the output of which passes through a static nonlinearity to predict the time-varying spike rate response (71, 72). The STRF, *h*(*x, u*) is as a linear weight matrix that is convolved with the logarithm of the stimulus spectrogram, *s*(*x, t*):

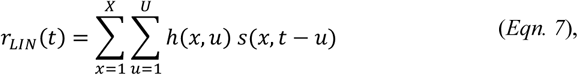

where *x* = 1…X are the frequency channels, *t* = 1…T is time, and *u* is the time lag of the convolution kernel. Taking the log of the stimulus spectrogram accounts for nonlinear gain in the cochlea. Free parameters in the weight matrix, *h*, indicate the gain applied to frequency channel *x* at time lag *u* to produce the predicted response. Positive values indicate components of the stimulus correlated with increased firing, and negative values indicate components correlated with decreased firing.

The output of the linear STRF is passed through a static nonlinear sigmoid function to account for spike threshold and saturation (34),

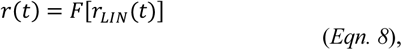

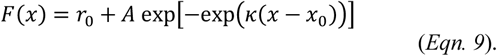

Free parameters here are *x*_*0*_, inflection point of the sigmoid, *r*_*0*_, spontaneous spike rate, *A*, maximum spike rate, and *κ*, the slope of the sigmoid.

We developed a modified LN STRF to account for stream-dependent changes in gain. The input spectrogram for each stream was scaled by a gain term that depended on stream identity (foreground or background) and trial phase (random or repeating). We refer to this model as the *phase*+*stream model*. The stimulus was modeled as the sum of two log spectrograms, computed separately for the foreground and background streams, *s*1 and *s*2, respectively. In the random phase, the total stimulus, *s*(*x, t*), was modeled as the linear sum of these two stimuli:

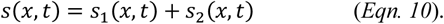

In the repeating phase, each stimulus was scaled by a repetition gain for the respective stream,

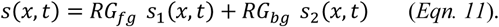

All model parameters were estimated by gradient descent (28, 34, 73). STRF parameters were initialized to have flat tuning (*i.e*., uniform initial values of *h*) and were iteratively updated using small steps in the direction that optimally reduced the mean squared error between the time-varying spike rate of the neuron and the model prediction. To maximize statistical power with the available data, the STRF was fit using both single- and dual-stream data. For single-stream trials, the second stimulus spectrogram was fixed at zero, *s*_2_(*x, t*) = 0, and a separate gain term was fit for those trials to prevent bias in estimates of *RG*_*fg*_ and *RG*_*bg*_. Measurements of prediction accuracy were obtained by 20-fold cross validation, in which a separate model was fit to 95% of the data then used to predict the remaining 5%. Fit and test data were taken from interleaved trials. This procedure was repeated 20 times with non-overlapping test sets, so that the final result was a prediction of the entire time-varying response. Prediction accuracy was then measured as the correlation coefficient (Pearson’s *r*) between the predicted and actual response. Standard error on prediction correlation was measured by jackknifing (74), and only units with prediction error significantly greater than zero were included in model comparisons (*p*<0.05, jackknife *t*-test).

To quantify effects of phase- and stream-dependent gain, we also fit models using the same data and fitting procedure, but where stream identity (phase-only model) or both phase and stream (baseline model) were shuffled in time. An improvement in prediction accuracy for a model with a non-shuffled over shuffled variable indicated a beneficial effect of the corresponding gain parameter on model performance, and thus of a stream-dependent change in sound encoding. Significant differences in model performance were assessed by a Wilcoxon rank sum test between prediction correlations for the set of units fit with each model.

## Author Contributions and Notes

D.S. and S.V.D. designed research, D.S. performed experiments, B.N.B. and S.V.D. wrote acquisition software, B.N.B, S.V.D., and D.S. analyzed data; and D.S. and S.V.D. wrote the paper. The authors declare no conflict of interest.

## Acknowledgments

The authors thank Dr. Josh H. McDermott for providing the code to generate the noise samples, and Zachary Schwartz and Henry Cooney for assistance with behavioral training and neurophysiological recordings. This study was supported by grants from the NIH (S.V.D., R00 DC 010439, R01 DC 014950) and the Hearing Health Foundation (B.N.B.).

## References

1. Griffiths TD, Warren JD (2004) What is an auditory object? Nat Rev Neurosci 5(11):887–92.

2. Bregman AS (1990) Auditory Scene Analysis: The Perceptual Organization of Sound (Cambridge, MA). MIT Press.

3. Carlyon RP (2004) How the brain separates sounds. Trends Cogn Sci 8(10):465–71.

4. Darwin CJ (1997) Auditory grouping. Trends Cogn Sci 1(9):327–33.

5. McDermott JH (2009) The cocktail party problem. Curr Biol 19(22):R1024–7.

6. Winkler I, Denham SL, Nelken I (2009) Modeling the auditory scene: predictive regularity representations and perceptual objects. Trends Cogn Sci 13(12):532–40.

7. Bregman AS (1978) Auditory streaming: competition among alternative organizations. Percept Psychophys 23(5):391–8.

8. Bregman AS, Ahad PA, Crum PA, O’Reilly J (2000) Effects of time intervals and tone durations on auditory stream segregation. Percept Psychophys 62(3):626–36.

9. Oberfeld D (2014) An objective measure of auditory stream segregation based on molecular psychophysics. Atten Percept Psychophys 76(3):829–51.

10. van Noorden L (1975) Temporal Coherence in the Perception of Tone Sequences. Dissertation (Technical University Eindhoven).

11. Akeroyd MA, Carlyon RP, Deeks JM (2005) Can dichotic pitches form two streams? J Acoust Soc Am 118(2):977–81.

12. Mesgarani N, Chang EF (2012) Selective cortical representation of attended speaker in multi-talker speech perception. Nature 485(7397):233–6.

13. Cusack R, Roberts B (2000) Effects of differences in timbre on sequential grouping. Percept Psychophys 62(5):1112–20.

14. Roberts B, Glasberg BR, Moore BCJ (2002) Primitive stream segregation of tone sequences without differences in fundamental frequency or passband. J Acoust Soc Am 112(5 Pt 1):2074–85.

15. Singh PG, Bregman AS (1997) The influence of different timbre attributes on the perceptual segregation of complex-tone sequences. J Acoust Soc Am 102(4):1943–52.

16. Micheyl C, Shamma SA, Oxenham AJ (2007) Hearing Out Repeating Elements in Randomly Varying Multitone Sequences: A Case of Streaming? BT - Hearing – From Sensory Processing to Perception. eds Kollmeier B, et al. (Springer Berlin Heidelberg, Berlin, Heidelberg), pp 267–274.

17. Shamma SA, Elhilali M, Micheyl C (2011) Temporal coherence and attention in auditory scene analysis. Trends Neurosci 34(3):114–23.

18. Elhilali M, Ma L, Micheyl C, Oxenham AJ, Shamma SA (2009) Temporal coherence in the perceptual organization and cortical representation of auditory scenes. Neuron 61(2):317–29.

19. Bendixen A, Denham SL, Gyimesi K, Winkler I (2010) Regular patterns stabilize auditory streams. J Acoust Soc Am 128(6):3658–66.

20. Szalárdy O, et al. (2014) The effects of rhythm and melody on auditory stream segregation. J Acoust Soc Am 135(3):1392–405.

21. Andreou L-V, Kashino M, Chait M (2011) The role of temporal regularity in auditory segregation. Hear Res 280(1–2):228–35.

22. Agus TR, Thorpe SJ, Pressnitzer D (2010) Rapid formation of robust auditory memories: insights from noise. Neuron 66(4):610–8.

23. McDermott JH, Wrobleski D, Oxenham AJ (2011) Recovering sound sources from embedded repetition. Proc Natl Acad Sci U S A 108(3):1188–93.

24. Pérez-González D, Malmierca MS (2014) Adaptation in the auditory system: an overview. Front Integr Neurosci 8:19.

25. Ulanovsky N, Las L, Nelken I (2003) Processing of low-probability sounds by cortical neurons. Nat Neurosci 6(4):391–8.

26. Nelken I (2014) Stimulus-specific adaptation and deviance detection in the auditory system: experiments and models. Biol Cybern 108(5):655–663.

27. Näätänen R (2001) The perception of speech sounds by the human brain as reflected by the mismatch negativity (MMN) and its magnetic equivalent (MMNm). Psychophysiology 38(1):1–21.

28. David S V. (2018) Incorporating behavioral and sensory context into spectro-temporal models of auditory encoding. Hear Res 360:107–123.

29. David S V, Fritz JB, Shamma SA (2012) Task reward structure shapes rapid receptive field plasticity in auditory cortex. Proc Natl Acad Sci U S A 109(6):2144–9.

30. Yin P, Fritz JB, Shamma SA (2010) Do ferrets perceive relative pitch? J Acoust Soc Am 127(3):1673–80.

31. Masutomi K, Barascud N, Kashino M, McDermott JH, Chait M (2015) Sound Segregation via Embedded Repetition Is Robust to Inattention. J Exp Psychol Hum Percept Perform. doi:10.1037/xhp0000147.

32. Vinje WE, Gallant JL (2000) Sparse coding and decorrelation in primary visual cortex during natural vision. Science 287(5456):1273–6.

33. Depireux DA, Simon JZ, Klein DJ, Shamma SA (2001) Spectrotemporal response field characterization with dynamic ripples in ferret primary auditory cortex. J … 85(3):1220–1234.

34. Thorson IL, Liénard J, David S V (2015) The Essential Complexity of Auditory Receptive Fields. PLoS Comput Biol 11(12):e1004628.

35. Bizley JK, Cohen YE (2013) The what, where and how of auditory-object perception. Nat Rev Neurosci 14(10):693–707.

36. Micheyl C, Tian B, Carlyon RP, Rauschecker JP (2005) Perceptual organization of tone sequences in the auditory cortex of awake macaques. Neuron 48(1):139–48.

37. Fishman YI, Reser DH, Arezzo JC, Steinschneider M (2001) Neural correlates of auditory stream segregation in primary auditory cortex of the awake monkey. Hear Res 151(1–2):167–187.

38. Teki S, Chait M, Kumar S, von Kriegstein K, Griffiths TD (2011) Brain bases for auditory stimulus-driven figure-ground segregation. J Neurosci 31(1):164–71.

39. Teki S, Chait M, Kumar S, Shamma SA, Griffiths TD (2013) Segregation of complex acoustic scenes based on temporal coherence. Elife 2:e00699.

40. Bar-Yosef O, Nelken I (2007) The effects of background noise on the neural responses to natural sounds in cat primary auditory cortex. Front Comput Neurosci 1:3.

41. Bar-Yosef O, Rotman Y, Nelken I (2002) Responses of neurons in cat primary auditory cortex to bird chirps: effects of temporal and spectral context. J Neurosci 22(19):8619–32.

42. Ulanovsky N, Las L, Farkas D, Nelken I (2004) Multiple time scales of adaptation in auditory cortex neurons. J Neurosci 24(46):10440–53.

43. Grill-Spector K, Henson R, Martin A (2006) Repetition and the brain: neural models of stimulus-specific effects. Trends Cogn Sci 10(1):14–23.

44. Ding N, Simon JZ (2012) Emergence of neural encoding of auditory objects while listening to competing speakers. Proc Natl Acad Sci U S A 109(29):11854–9.

45. Mesgarani N, David S V, Fritz JB, Shamma SA (2009) Influence of context and behavior on stimulus reconstruction from neural activity in primary auditory cortex. J Neurophysiol 102(6):3329–39.

46. May PJC, Tiitinen H (2010) Mismatch negativity (MMN), the deviance-elicited auditory deflection, explained. Psychophysiology 47(1):66–122.

47. Anderson LA, Christianson GB, Linden JF (2009) Stimulus-specific adaptation occurs in the auditory thalamus. J Neurosci 29(22):7359–63.

48. Malmierca MS, et al. (2009) Stimulus-Specific Adaptation in the Inferior Colliculus of the Anesthetized Rat. J Neurosci 29(17):5483–5493.

49. Antunes FM, Nelken I, Covey E, Malmierca MS (2010) Stimulus-Specific Adaptation in the Auditory Thalamus of the Anesthetized Rat. PLoS One 5(11):e14071.

50. Nelken I, Ulanovsky N (2007) Mismatch Negativity and Stimulus-Specific Adaptation in Animal Models. J Psychophysiol 21(3–4):214–223.

51. Malmierca MS, Anderson LA, Antunes FM (2015) The cortical modulation of stimulus-specific adaptation in the auditory midbrain and thalamus: a potential neuronal correlate for predictive coding. Front Syst Neurosci 9:19.

52. Natan RG, et al. (2015) Complementary control of sensory adaptation by two types of cortical interneurons. Elife 4. doi:10.7554/eLife.09868.

53. Yarden TS, Nelken I (2017) Stimulus-specific adaptation in a recurrent network model of primary auditory cortex. PLOS Comput Biol 13(3):e1005437.

54. Ayala YA, Malmierca MS (2013) Stimulus-specific adaptation and deviance detection in the inferior colliculus. Front Neural Circuits 6:89.

55. Taaseh N, Yaron A, Nelken I (2011) Stimulus-specific adaptation and deviance detection in the rat auditory cortex. PLoS One 6(8):e23369.

56. Darwin C, Carlyon RP (1995) Auditory grouping: The Handbook of Perception and Cognition (ed Moore BCJ, Academic, New York). Vol 6.

57. Yost WA, Popper AN, Fay RR eds. (2007) Auditory Perception of Sound Sources (Springer US, Boston, MA) doi:10.1007/978-0-387-71305-2.

58. Fay R (2008) Sound source perception and stream segregation in nonhuman vertebrate animals. Auditory Perception of Sound Sources., eds Yost W, Fay R, Popper A (Springer; New York), pp 307–323.

59. Bee MA, Micheyl C (2008) The cocktail party problem: what is it? How can it be solved? And why should animal behaviorists study it? J Comp Psychol 122(3):235–51.

60. Ma L, Micheyl C, Yin P, Oxenham AJ, Shamma SA (2010) Behavioral measures of auditory streaming in ferrets (Mustela putorius). J Comp Psychol 124(3):317–30.

61. Cohen MR, Maunsell JHR (2009) Attention improves performance primarily by reducing interneuronal correlations. Nat Neurosci 12(12):1594–600.

62. Sundberg K a, Mitchell JF, Reynolds JH (2009) Spatial attention modulates center-surround interactions in macaque visual area v4. Neuron 61(6):952–63.

63. Reynolds JH, Chelazzi L (2004) Attentional modulation of visual processing. Annu Rev Neurosci 27:611–47.

64. Barascud N, Pearce MT, Griffiths TD, Friston KJ, Chait M (2016) Brain responses in humans reveal ideal observer-like sensitivity to complex acoustic patterns. Proc Natl Acad Sci 113(5):E616–E625.

65. Cherry EC (1953) Some Experiments on the Recognition of Speech, with One and with Two Ears. J Acoust Soc Am 25(5):975–979.

66. Slee SJ, David S V. (2015) Rapid task-related plasticity of spectro-temporal receptive fields in the auditory midbrain. J Neurosci 35(38):13090–102.

67. Bizley JK, Nodal FR, Nelken I, King AJ (2005) Functional organization of ferret auditory cortex. Cereb Cortex 15(10):1637–53.

68. Englitz B, David SV V, Sorenson MDD, Shamma SA (2013) MANTA - An open-source, high density electrophysiology recording suite for MATLAB. Front Neural Circuits 7:69.

69. David S V, Mesgarani N, Fritz JB, Shamma SA (2009) Rapid synaptic depression explains nonlinear modulation of spectrotemporal tuning in primary auditory cortex by natural stimuli. J Neurosci 29(11):3374–86.

70. Hoffman MD, Gelman A (2011) The No-U-Turn Sampler: Adaptively Setting Path Lengths in Hamiltonian Monte Carlo. Available at: http://arxiv.org/abs/1111.4246 [Accessed February 10, 2019].

71. Aertsen AM, Johannesma PI (1981) The spectro-temporal receptive field. A functional characteristic of auditory neurons. Biol Cybern 42(2):133–43.

72. deCharms RC, Blake DT, Merzenich MM (1998) Optimizing sound features for cortical neurons. Science 280(5368):1439–43.

73. Byrd RH, Lu P, Nocedal J, Zhu C (1995) A Limited Memory Algorithm for Bound Constrained Optimization. SIAM J Sci Comput 16(5):1190–1208.

74. Efron B, Tibshirani R (1986) Bootstrap Methods for Standard Errors, Confidence Intervals, and Other Measures of Statistical Accuracy. Stat Sci 1(1):54–75.

75. Fox J (1997) Applied regression analysis, linear models, and related methods. - PsycNET Available at: https://psycnet.apa.org/record/1997-08857-000 [Accessed August 16, 2019].

